# Adaptation of a methanogen to the constructed environment

**DOI:** 10.1101/2024.03.06.583737

**Authors:** Satoshi Kawaichi, Rhitu Kotoky, Jacek Fiutowski, Amelia-Elena Rotaru

## Abstract

Due to unique genomic adaptations, Methanococcus maripaludis Mic1c10 exhibits severe corrosive behavior when in direct contact with Fe0. These adaptations are linked to attachment and effective growth on constructed surfaces. One such adaptation is that of a specific [NiFe]-hydrogenase that may anchor on the cell surface via glycosyl-glycosyl interactions to receive Fe0-electrons directly. Such an evolutionary response to constructed environments requires us to rethink methane cycling in human-altered ecosystems.

---

Iron (Fe^0^) corrosion in anoxic environments can be caused by methanogenic archaea, working alone or in conjunction with bacteria. Certain strains of *Methanococcus maripaludis* are particularly corrosive and are found to be associated with infrastructure corrosion globally^1^.

Previous studies on mutant strains of a non-corrosive *M. maripaludis* (S2) indicated that biological corrosion occurs due to the release of free enzymes from moribund cells during the late-stationary phase^2^. These enzymes, such as extracellular hydrogenases, can catalyze electron uptake from Fe^0^ and hydrogen evolution^3^. The hydrogen is then used as an electron donor by active cells of *M. maripaludis*^2^.

Further studies have suggested that free enzymes released by a severely corrosive *M. maripaludis* (OS7), are the primary cause for its corrosive capabilities^4^. The strain contains a conserved yet genetically unstable genetic island, the Microbiologically Influenced Corrosion Island (MIC Island), with genes for two subunits of a [NiFe]-hydrogenase and the Tat transmembrane protein secretion system, believed to export the hydrogenases outside the cell. Researchers found the [NiFe]-hydrogenases in spent cell filtrates and determined that the spontaneous deletion of the MIC-island in a mutant strain (OS7mut1) resulted in a loss of the ability to corrode. This highlights the crucial role this island plays in corrosion^4^.

Moreover, when a segment of the MIC-island that included the [NiFe]-hydrogenases and the Tat-system was transplanted from the corrosive strain OS7 to a non-corrosive strain JJ, the resulting mutant exhibited some corrosive properties^5^, albeit an order of magnitude lower than corrosive OS7. This suggests there might be additional factors required for full corrosive potency. Complementing this insight, Holten *et al*.^5^ showed that glycosylated S-layer proteins facilitate the attachment of *M. maripaludis* to surfaces, including Fe^0^.

While previous studies indicate that corrosive *M. maripaludis* rely on free enzymes to access Fe^0^-electrons, free enzymes are costly as the producer makes them readily available to the entire community, including ‘cheater’ populations^6^. The cost is particularly high when such enzymes produce a highly diffusible product like hydrogen, which becomes a “public good”^6^. Therefore, retaining the enzymes close to the producer cells can provide a competitive advantage against ‘cheater’ populations.

Thus, we hypothesize that highly corrosive strains of *M. maripaludis* keep their extracellular enzymes close to the cell surface necessitating direct contact to the Fe^0^-surface. This enables effective electron uptake from Fe^0^ and corrosion. To test this hypothesis, we conducted physiological experiments and compared the genome of a corrosive strain *-* Mic1c10^7^ - with three corrosive^1,4,8^ and three non-corrosive^9,10^ *M. maripaludis*. Our results show that corrosive cells directly interact with the metal surface to retrieve electrons, likely through [NiFe]-hydrogenases attached to cell surface proteins via glycosyl-glycosyl interactions.

We found that physical contact was crucial for corrosion by *M. maripaludis* strain Mic1c10. When Mic1c10 interacted with a carbon steel surface, it created a distinctive black film and caused permanent surface alterations to the material surface, which were not observed in cell-free controls (Fig. 1a-1c). With its small cell size (∼500 nm diameter) and high surface-to-volume ratio, it could effectively attach, colonize, and corrode the metal surface (Fig. 1d, Table 1SM). The number of Mic1c10 cells doubled every 2.5 days until they reached a plateau after two weeks. The cell count remained steady for another four weeks (Fig. 1e).

**Figure 1.**
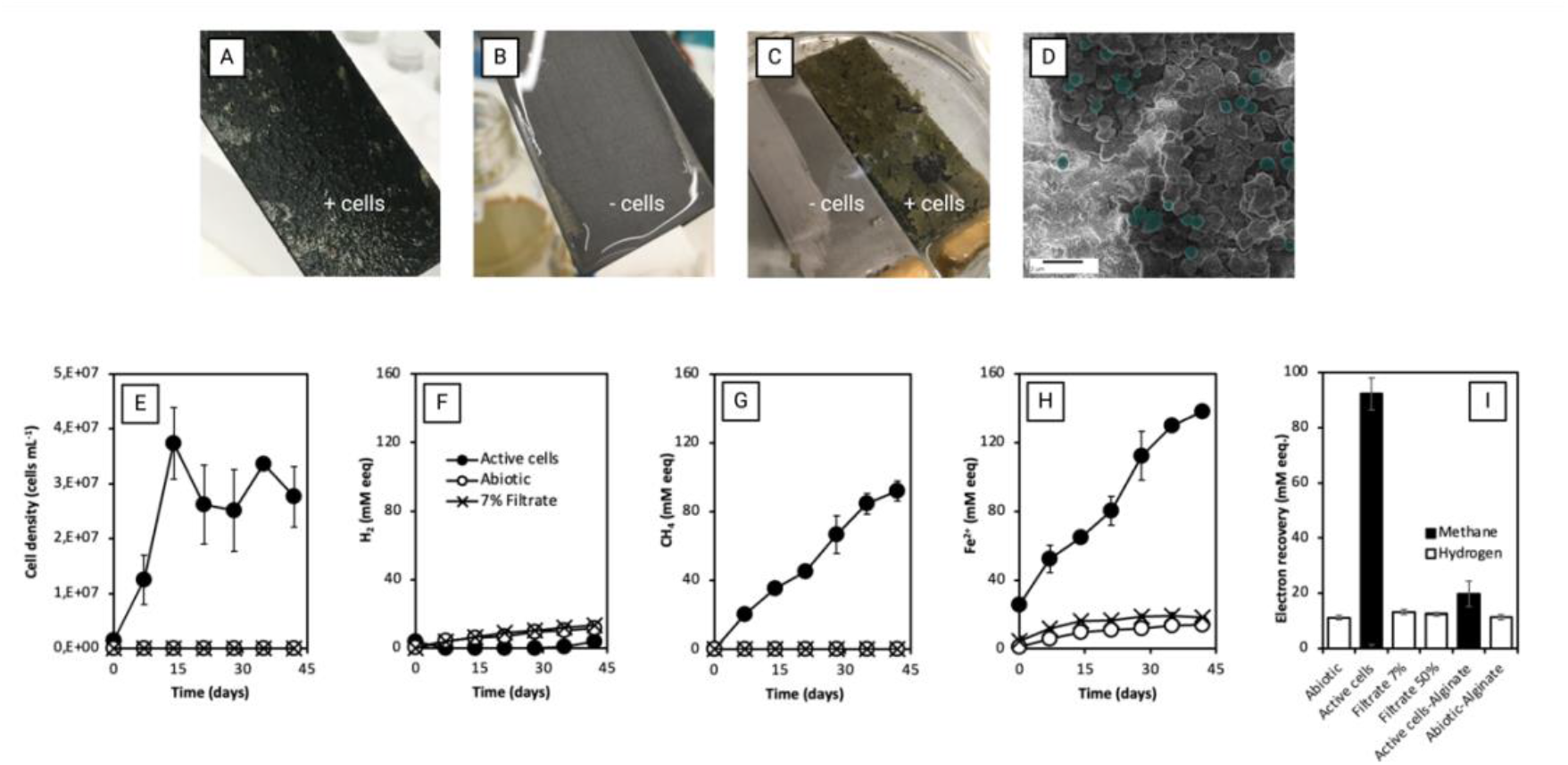
Corrosion by *M. maripaludis* Mic1c10. (A) Cells grown on carbon steel compared to (B) control without cells, both before, and (C) after acid-washing to remove loosely adhered material, highlighting corrosion. (D) HIM image showcasing Mic1c10 colonization on Fe^0^-foil. (E) Cell quantification using DAPI staining after vigorous shacking to detach cells from Fe^0^-foil. (F-H) Measurement of corrosion products: hydrogen, methane, and ferrous iron (in mM electron equivalents). (I) Electron recovery comparison across treatments. All tests were performed with several biological replicates (n≥3).

Methane and Fe^2+^ were the primary products of Mic1c10’s corrosive metabolism throughout the entire six-week period (Fig. 1f, 1g, 1h). This was due to the methanogenic metabolism of Mic1c10, which drove the reduction of CO_2_ to methane and ferrous iron (Fe^2+^) via the MIC reaction (4Fe^0^ + 5HCO_3_ ^-^ + 5H^+^ → CH_4_ + 4FeCO_3_ + 3H_2_O). On the contrary, abiotic controls accumulated hydrogen and Fe^2+^ according to an abiotic reaction (Fe^0^ + HCO_3_ ^-^ + H^+^ → FeCO_3_ + H_2_).

Spent filtrate is supposed to contain enzymes that promote Fe^0^ corrosion even in the absence of active cells^2,4^. However, the addition of spent filtrate from Mic1c10 to fresh media with Fe^0^ resulted in negligible H_2_- and formate-buildup (0.16 ± 0.04 mM), which could explain less than 3% of the methane formed by cells (Fig. 1, Figure 1SM). Furthermore, spent filtrate did not promote accumulation of Fe^2+^ a direct product of Fe^0^-oxidation above abiotic controls (p=0.45, n>3), clearly showing that Mic1c10’s filtrate lacked intrinsic corrosive properties whereas Mic1c10 were severely corrosive. These findings are in contrast to earlier reports^2,4^, that highlighted the corrosive influence of free enzymes. We thus directed our focus to cell-bound elements as potential primary agents for severe corrosion, since soluble extracellular constituents in spent filtrate had minimal impact.

To investigate, we applied an alginate hydrogel coating on the Fe^0^ surface, which allows molecular diffusion of hydrogen while blocking cell contact with the metal. This setup was aimed to dissect the role of physical contact in the corrosion process. We observed that cells interacting with alginate-coated-Fe^0^ exhibited a fivefold decrease in methane production compared to those in direct contact with uncoated Fe^0^. (Fig. 1i, Fig. 2SM). This suggests that Mic1c10 was inefficient in extracting electrons from Fe^0^ without direct contact, emphasizing the critical role of physical interaction in the microbial corrosion mechanism.

**Figure 2.**
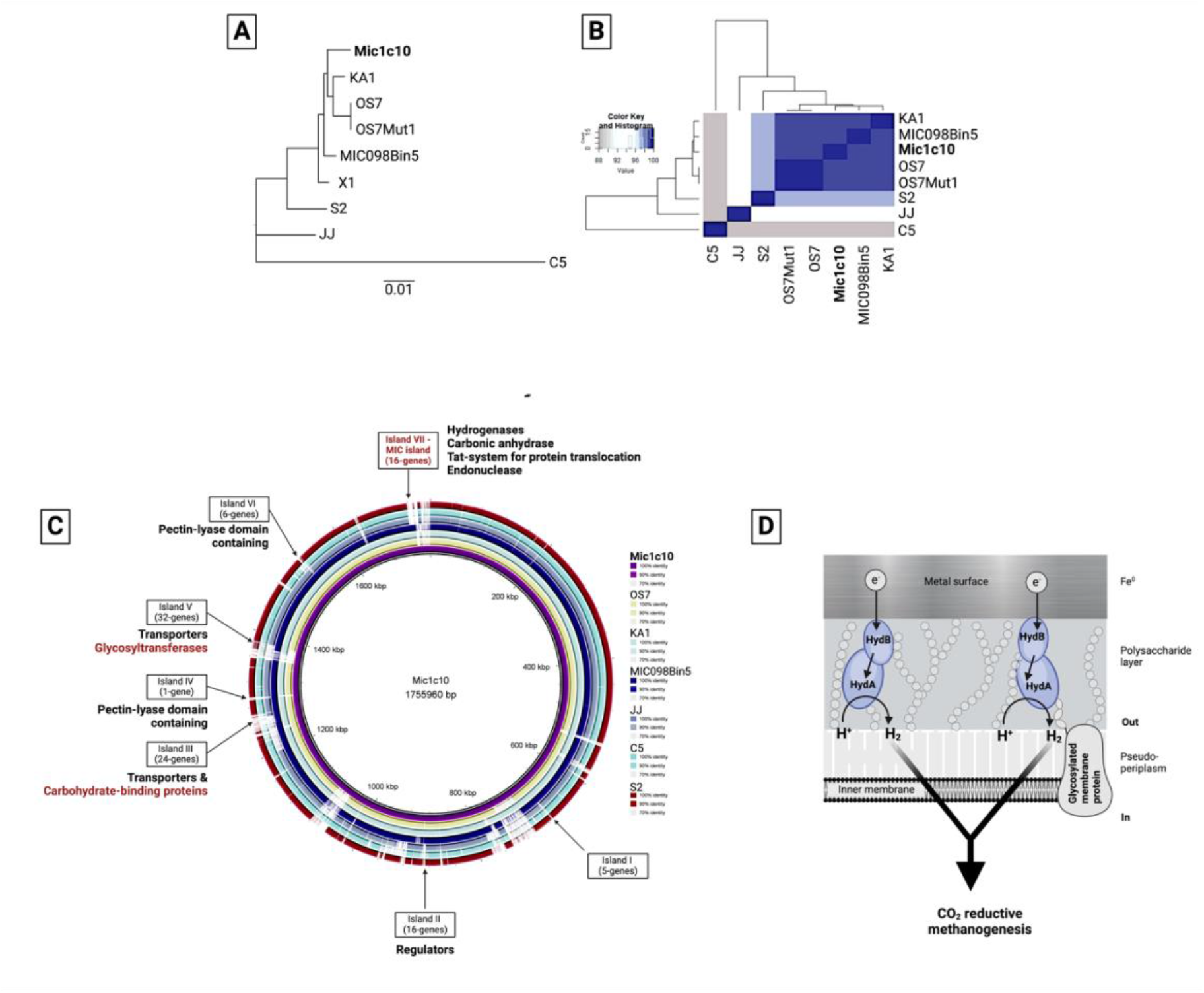
Comparative genomics of *Methanococcus maripaludis* Mic1c10 against corrosive and non-corrosive strains. (A) Core genome MLST similarity tree of 453 concatenated genes (B) average nucleotide identity and (C) whole genome alignments for identification of divergent genetic regions between strains. (D) Predicted model of electron uptake from the Fe^0^-surface.

The next objective was to understand what makes Mic1c10 cells efficiently extract electrons from Fe0, leading to severe corrosion. We hypothesized that Mic1c10 uses surface-anchored hydrogenases to facilitate electron uptake from Fe^0^ (Fig. 2). To test this hypothesis and understand the genetic basis of its severely corrosive behavior, we conducted a comprehensive comparative genomic analysis. First, the genome of Mic1c10 was sequenced and compared to that of non-corrosive (JJ^9,10^, C5 and S2^10^) along with corrosive *M. maripaludis* strains (KA1^8^ and OS7^4^) and a nearly complete metagenome-assembled genome (MAG) of a strain linked to global infrastructure corrosion^1^.

Our analysis revealed a close evolutionary link among corrosive strains based on 453 genes, while non-corrosive strains showed a marked divergence from corrosive strains (Fig. 2a). ANI values supported these findings with Mic1c10 sharing high average nucleotide identity (ANI) to other corrosive strains (≥98.5%) and lower ANI to non-corrosive strains (<97%). Further examination of the genomic differences between corrosive and non-corrosive strains led to the identification of seven genomic regions unique to the corrosive strains (Fig. 2c). These regions encoded functions that appear critical for the corrosive capabilities of the strains, suggesting that the ability to cause severe corrosion is a genetically-supported trait.

Mic1c10, like all other corrosive strains, presented a specific MIC island that contained the two subunits of a bidirectional [NiFe]-hydrogenase^4^. Our *in-silico* predictions suggested that these subunits formed a dimer (Fig. 3SM) with external protrusions that could be glycosylated. Together, the two subunits had nine N-x-T/S motifs specifying glycosylation sites (Fig. 3SM, Table S3). Interestingly, the [NiFe]-hydrogenase of Mic1c10 had four additional glycosylation sites than those of other corrosive strains, suggesting that it may become more adhesive if fully glycosylated. A tight anchorage of this enzyme could also explain why we could not observe hydrogenase activity in Mic1c10’s cell filtrates (Fig.1).

We suspect that glycosylated [NiFe]-hydrogenases could be anchored onto glycosylated S-layer proteins previously reported in this methanogen^5^. For this, the cells require glycosyl transferases such as AglB to glycosylate the hydrogenases and the S-layer proteins^5^. While AglB is not unique to corrosive strains, seven glycosyl transferases and four additional enzyme possibly catalyzing the biosynthesis of unusual sugars and deoxy-sugars for cell wall synthesis^11^ and S-layer glycosylations^12^, were encoded in a genomic-region no. V which is unique to corrosive strains (Fig. 2c, Table S2). These results indicate that glycosylated S-layer proteins may anchor glycosylated [NiFe]-hydrogenases on the cell’s exterior (Fig. 2d). This supports our hypothesis that Mic1c10 uses surface-anchored hydrogenases to facilitate electron uptake from Fe^0^ (Fig. 2).

Moreover, the other genomic islands specific to Fe^0^-corrroders, contained various genes involved in gene expression regulation (island no. II), transport of metals, siderrophores, aminoacids and vitamins (especially island no. III), adhession and biofilm formation (islands III and V) indicating an adaptation to interactions with metalic surfaces. Interestingly, the high similarity of Mic1c10’s genome and other corrosive *M. maripaludis* genomes to an environmental variant responsible for corroding global infrastructure indicates a shared evolutionary adaptation to the constructed environment and underscores the ecological relevance of our findings.

In conclusion, we found that Mic1c10 requires direct contact between cells and metal to uptake electrons, challenging the traditional view of free enzyme’s role in corrosion by *Methanococcus maripaludis*. We identified a ‘corrosive core genome’ consisting of seven genomic regions containing [NiFe]-hydrogenases, regulatory elements and various enzymes involved in protein glycosylation. Glycosylation could explain how hydrogenases remain anchored on the cell surface. The intraspecies diversity of *M. maripaludis*, suggests a high adaptive potential. Minor genetic variations, such as additional glycosylation sites on a hydrogenase, affect its phenotypic behavior. Our findings have implications for developing strategies to control MIC such as designing glycosylation inhibitors or engineering microbial strains with enhanced glycosylation capabilities for biotechnology.

## Acknowledgements

This grant is a contribution to a Sapere Aude FNU grant awarded to AER (grant number 4181-00203). Other support was by the Novo Nordisk Foundation grant NNF21OC0067353, a Danish Research Council grant DFF 1026-00159, and a European Research Council ERC-CoG grant 101045149 awarded to AER. JF is supported by UFM 5229-00010B, NanoChem for National Research Infrastructure.

We would like to thank Arkadiusz Goszczak for SEM-EDX, Oona Snoeyenbos-West for valuable discussions and Mon Oo Yee, Paola Palacios, Lasse Ørum Smidt, and Kanami Kawaichi for lab assistance.

## Materials and Methods [Online only]

### Strains and batch culture conditions

*M. maripaludis* Mic1c10 (= NBRC 105639) was routinely cultured on a modified NBRC medium 927 with reduced amount of Fe^0^ granules to 100 g/L, and addition of 1 mg/L resazurin as an oxygen indicator. The medium and stock solutions were prepared anaerobically by degassing with N_2_/CO_2_ (80/20). All culture experiments were carried out in 120 mL serum bottles containing 50 mL culture, a gas phase of N_2_/CO_2_ (80/20), and 5.0 g Fe^0^ granules (1 to 2 mm in diameter, 99.98% purity metal basis; Alpha Aesar, Ward Hill, MA) at 37°C unless otherwise mentioned. Any culture with sign of oxygen contamination was removed from the experiments. Amount of CH_4_, H_2_, and Fe^2+^, and the cell density were measured periodically (see below).

### Preparation of cell-free spent culture medium (SCM)

SCM was prepared by filtering a one-month-old culture (late exponential growth phase) on Fe^0^ through two stacked syringe filters (cellulose acetate membrane, 0.2 μm pore size; GVS Filter Technology), essentially as described by Deutzmann *et al*^2^. To initiate the experiment, 3 and 25 mL of SCM was added anoxically to 40 and 25 mL fresh medium, respectively, with 5.0 g Fe^0^ granules.

### Encapsulation of Fe^0^ with alginate hydrogel

For the alginate experiment, 5 mL of 1% (w/v) alginic acid sodium salt solution was added to a 120 mL serum bottle with 5.0 g iron granules. Then the bottle was degassed with N_2_/CO_2_ (80/20) for 10 minutes and autoclaved. Ten milliliters of 1% (w/v) CaCl_2_ solution was slowly added to the bottle to overlay on the alginic acid solution, and the bottle was allowed to stand still overnight to solidify the hydrogel. The remaining CaCl_2_ solution was removed, and the solidified hydrogel was washed twice with 10 mL of fresh medium. Finally, 45 mL of fresh medium was added to the bottle. To initiate the experiment, 5 mL of well-grown culture was added to the bottle.

### Analytical procedures

CH_4_ and H_2_ were measured using a gas chromatograph (GC) (Trace 1300, Thermo Scientific) equipped with a thermal conductivity detector (TCD). The injector was operated at 150°C and the TCD at 200°C with 1.0 mL/min reference gas flow. The oven temperature was constant at 70°C. Gas chromatography analyses used a TG-BOND Msieve 5A column (30-m length, 0.53-mm i.d., and 20-μm film thickness, Thermo Scientific) with argon carrier gas at 25 mL/min. The GC was controlled and automated by the Chromeleon software (Dionex, Version 7).

The amount of Fe^2+^ was measured using a modified ferrozine method^13^. A 100 μL sample was acidified with 900μL of 0.5M HCl, 10 μL of the mixture was added to 2 mL of 0.2g/L 3-(2-pyridil)-5,6-bis(4-phenylsulphonate)-1,2,4-triazin (ferrozine) solution in 50 mM HEPES at pH 7.0, and the absorbance at 562 nm was measured.

Cell growth was monitored by the direct cell count of 4’,6-diamidino-2-phenylindole (DAPI) stained cells using an epifluorescent microscope. Mineral precipitates in the sampled culture were dissolved using oxalate before DAPI-staining. A 500 μL sample was collected anaerobically and an equal volume of marine TPE solution (100 mM Tris pH 7.0, 10 mM EDTA, and 300 mM sodium phosphate) was added without significantly changing the osmolality. Filter-sterilized oxalate solution (200 mM ammonium oxalate, and 120 mM oxalic acid) was added until the culture turns yellow. Cells in the mixture were collected on a membrane filter (pore size; 0.2 μm, Millipore). The effect of the oxalate treatment on the microbial cell counts was evaluated by counting cells from H_2_ grown culture with or without the treatment and confirmed to be negligible (data not shown).

### Whole-cell cyclic voltammetry on carbon electrode

A two-chambered reactor with 300 mL volume (150 mL × 2) was filled with 200 mL modified NBRC medium 927 without Fe^0^ as the electrolyte. Graphite electrodes (75 × 25 × 10 mm) were used as the working and counter electrodes. The electrode was fixed to a titanium wire using a conductive epoxy (EPO-TEK H20E, EPOXY TECHNOLOGY), and then the connection part was covered with non-conductive epoxy (EPO-TEK 730, EPOXY TECHNOLOGY). A leak-free Ag/AgCl (3.4 M KCl) electrode was used as the reference electrode. Cyclic voltammetry was performed using a MultiEmStat3+ potentiostat (PalmSens BV, Houten, The Netherlands) with a scan rage of -1.2 to 0.2 V (vs. Ag/AgCl 3.4 M KCl) and a scan rate of 1 mV/sec.

### Helium-ion microscopy

Strain Mic1c10 was incubated in 5 mL of modified NBRC medium 927 with an Fe^0^ foil (99.5%, 5 x 5 x 0.127 mm, Thermo Scientific Chemicals) in a 20 mL serum vial for 10 days. The foil was then fixed using 2% glutaraldehyde in marine TPE solution for 72 hours. The specimen was washed three times with the marine TPE solution, dehydrated in different ethanol dilutions (50, 70, 80, 90, 95, and 100% x 3 times, 10 minutes each) and hexamethyldisilazane (30 sec.), and dried under a continuous flow of N_2_. All procedures were done anoxically in the serum vial.

Microbial cells were imaged using a Zeiss ORION NanoFAB Helium Ion Microscope with SE detection (Zeiss, Germany). He+ imaging was performed at 25 keV beam energy, with a probe current ranging from 0.08 to 0.01 pA, and a scan dwell time of 1 μs. Charge compensation was applied, if necessary, using a low-energy electron beam, a flood gun with 433 eV. Sample working distance is 8.7 mm.

### Genome sequencing and taxonomic analysis

The whole genome of the strain Mic1c10 was sequenced through Illumina MiSeq paired-end library technique by FASMAC Co. Ltd. (Japan). The genome sequence was assembled using SPAdes (v. 3.15.3)^14^ with iterated k-value 21, 33, 55, 77, 99, and 127. All the scaffolds were annotated using Rapid prokaryotic genome annotation system (PROKKA)^15^ and BlastKOALA (ver. 2.2)^16^. To resolve the taxonomy of Mic1c10, the sequence of the isolate was compared to the Type Strain Genome Server (TYGS) based on 16s rDNA and whole genome sequence.

### Phylogenomic analysis of Mic1c10

The genome of Mic1c10 was compared with all complete genomes of *Methanococcus maripaludis* (refseq = 9) for the Average nucleotide identity (ANI). The ANI values (all vs all) were computed using an alignment-free fastANI v1.33^17^ algorithm to identify the closest reference genome by orthologous mappings and alignment identity estimates. The similarity matrix was used to generate a heatmap using ggplot2 in R version 3.6.3^18^.

All the complete genomes of *Methanococcus maripaludis* (refseq: 9 complete) including Mic1c10 were annotated using PROKKA and were used to construct the pangenome. All paralogous genes were rejected from the pan-genome allele and each orthologous cluster consisted of a protein family (a gene) with 95% sequence identity. The core genome multilocus sequence typing (MLST) analysis was done for all the core genome sequences (cgMLST) retrieved in the pangenome analysis and a core-genome phylogenetic tree was prepared using PHYLIP^19^ and FigTree v1.4.4 (http://tree.bio.ed.ac.uk/software/figtree/). The selected genomic features were compared using MAUVE (Multiple alignment of Conserved genomic sequence with rearrangement) ^20^ and BRIG (Blast Ring Image Generator)^21^ to identify uniques genes and genomic regions predictively related to corrosion.

### Identification of MIC (Microbe Induced Corrosion) island

The annotated features of the complete genomes were compared to identify the presence of the MIC island and further analyzed and visualized through alignment using MAUVE^20^ and BRIG^21^. For genomic feature alignment, the sequence of *M. maripaludis* OS7 (First report of MIC island sequence) was considered as reference genome and Genomic alignments were conducted using the progressive mauve tool with default settings. The selected genomes of corrosive and non-corrosive *M. maripaludis* strains were compared with BRIG to identify the pattern of unique or distinct genomic regions among the corrosive and non-corrosive strains.

### Other genome/protein analyses

The genes encoded in the seven genomic islands found in the corrosive strains were analyzed using InterPro^22^, ColabFold^23^, PEPPI^24^, and NetNGlyc 1.0^25^ to predict the protein localization, protein structure, protein-protein interactions, and potential glycosylation sites.

**Figures**

Composite figures were compiled using Biorender (https://www.biorender.com)

## Notes

### Competing Interest Statement

The authors have declared no competing interest.

