## Supplementary file for "Adaptation of a methanogen to the constructed environment"

**Supplementary Table S1. Tafel parameters of carbon-steel sheets incubated for 2 weeks with intact or heat-killed cells of strain Mic1c10 *.** The average and standard deviation of triplicate reactors for each condition are shown.

| | **Mic1c10 intact cells** | **Mic1c10 heat-killed cells** |
|---|---|---|
| **$J_{corr}$ (μA/cm$^2$)** | 186.7 ± 138.1 | 1.3 ± 1.0 |
| **$E_{corr}$ (mV)** | -594.3 ± 13.4 | -657.9 ± 17.0 |
| **$b_a$ (mV)** | 177.3 ± 54.3 | 44.7 ± 12.3 |
| **$b_c$ (mV)** | 625.8 ± 167.0 | 86.0 ± 23.7 |

* Electrochemical measurements of corrosion activity were conducted using a three-electrode single-cambered electrochemical reactor with a volume of 1 L filled with 500 ml culture medium. A carbon felt and an Ag/AgCl (saturated KCl) were used as counter and reference electrodes, respectively. The working electrode was prepared by polishing a mild steel coupon (25 x 75 x 1 mm) to a 1,200-grit finish. Subsequently, the coupon was fixed to a titanium wire with conductive epoxy (EPO-TEK H20E, EPOXY TECHNOLOGY). Following drying, the connection part was completely covered with non-conductive epoxy (EPO-TEK 730, EPOXY TECHNOLOGY), and then the working electrode was sterilized

with 96% (v/v) ethanol. The reactor was maintained under open-circuit conditions at room temperature to facilitate corrosion measurement. Linear sweep voltammetry (LSV) was periodically performed using an AMEL2553 potentiostat/galvanostat (Amel Electrochemistry, Milan, Italy) with a scan rate of 0.167 mV/sec. The initial scan range was established from -800mV to -500mV and adjusted to hold the corrosion potential within the scan range. The resultant current-potential (I-V) curve was analyzed using a Tafel fitting tool in CView ver. 3.5a (Scribner Associates Inc. NC, USA). This facilitated the calculation of corrosion parameters, including corrosion current ($I_{corr}$) and corrosion potential ($E_{corr}$). The corrosion current density ($J_{corr}$) was further derived from the $I_{corr}$ and the known surface area of the steel coupon (39.5 $cm^2$).

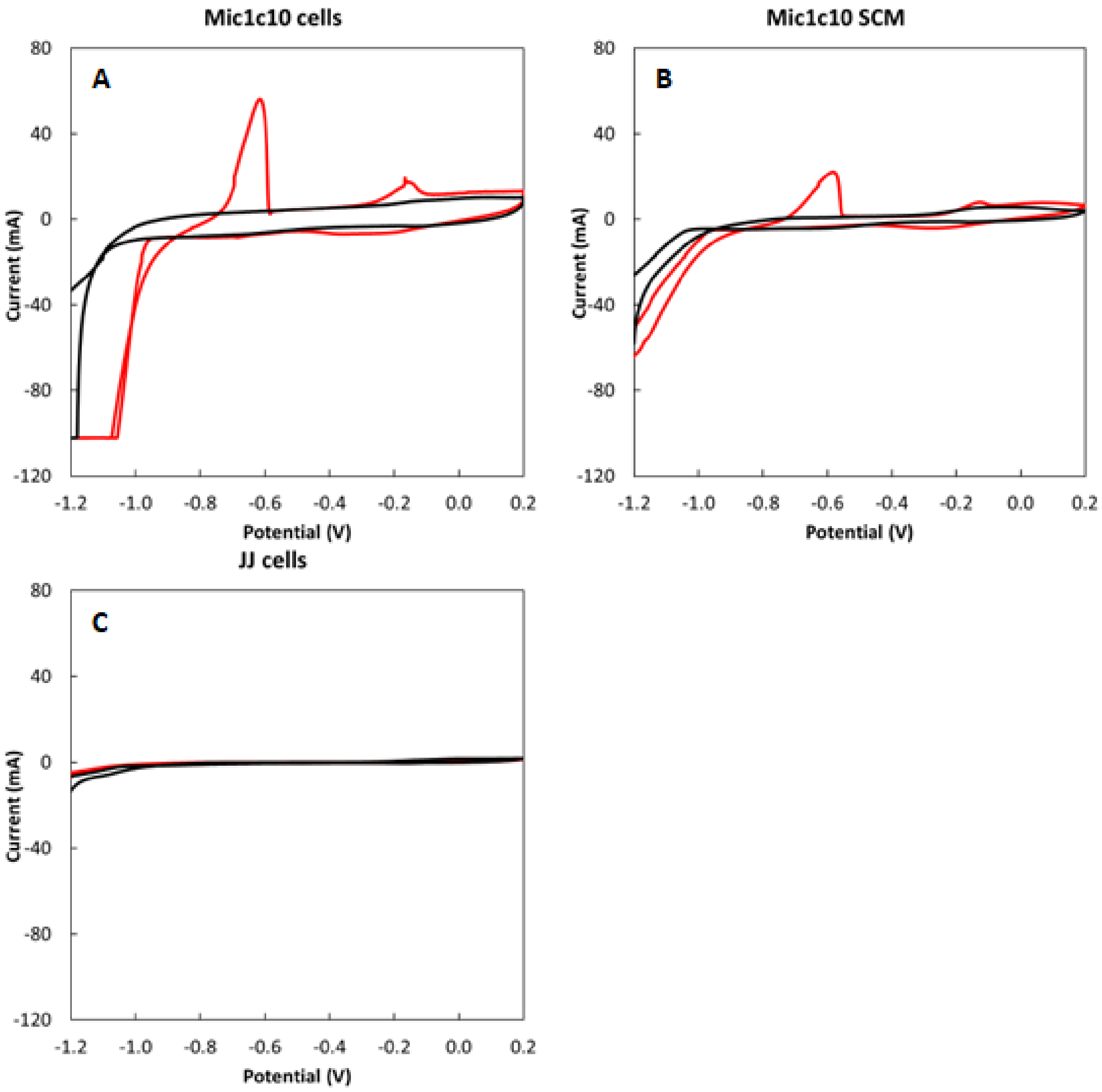


**Supplementary Figure 1. Cyclic voltammograms of the culture of strain Mic1c10 grown on $Fe^0$ (A) and its SCM (B) and that of the non-corrosive strain JJ cells (C) (red traces).** Voltammograms of the blank medium in each reactor are shown in black traces. The scan rate of the cyclic voltammetry (CV) was 1 mV/sec. A two-chambered reactor with 300 mL volume (150 mL × 2) was filled with 200 mL modified NBRC medium 927 without $Fe^0$ granules, as the electrolyte. Graphite electrodes (75 × 25 × 10 mm) were used as the working and counter electrodes. The electrode was fixed to a titanium wire using the conductive epoxy, and then the connection part was covered with the non-conductive epoxy. A leak-free Ag/AgCl (3.4 M KCl) electrode was used as the reference electrode. CV was performed using a MultiEmStat3+ potentiostat (PalmSens BV, Houten, The Netherlands) with a scan range of -1.2 to 0.2 V (vs. Ag/AgCl 3.4 M KCl). CV on a blank medium was measured first, and then 50% of the medium was replaced with the inoculum (Mic1c10, SCM, or JJ).

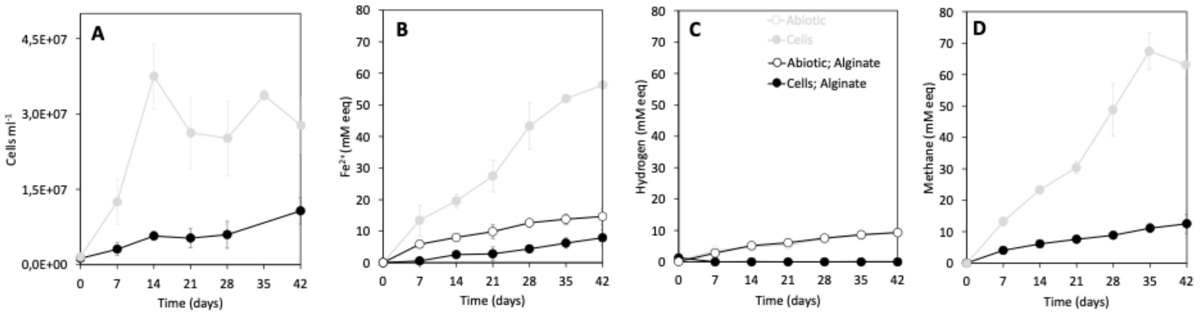


**Supplementary Figure 2. Incubations with $Fe^0$ coated with alginate.** (A) Cells grown on $H_2$ that crossed the alginate barrier (black circles) versus cells grown on $Fe^0$-directly (gray circles); (B) Formation of ferrous iron ($Fe^{2+}$), (C) hydrogen and (D) methane by cells grown on $Fe^0$ (gray circles), or on alginate-coated $Fe^0$ (black circles) versus abiotic $Fe^{2+}$ buildup on alginate-coated $Fe^0$ (empty circles). All incubations were carried out at a minimum in triplicate (n≥3).

| Position | Gene number in Mic1c10 | Annotation | Transmembrane helix: Phobius TM | Transmembrane helix: TMHMM | Signal peptide | N-X-S/T motif |
|---|---|---|---|---|---|---|
| | 07250 | hypothetical protein | 3 | 3 | No | 0 |
| | 07260 | Lysine exporter LysO (IPR005642) | 5 | 5 | Yes | 1 |
| I | 07270 | hypothetical protein | 0 | 0 | Yes | 2 |
| | 07280 | hypothetical protein | 0 | 0 | No | 0 |
| | 07290 | non-cytoplasmic hypothetical protein | 0 | 0 | Yes | 3 |
| | 09440 | hypothetical/ ClpP/crotonase - domain | 0 | 1 | No | 1 |
| | 09450 | hypothetical/ membrane bound | 3 | 3 | No | 1 |
| | 09460 | hypothetical/ membrane bound | 3 | 3 | No | 0 |
| | 09470 | Bax inhibitor 1 like (IPR010539) | 0 | 7 | No | 3 |
| | 09480 | hypothetical protein | 0 | 0 | No | 0 |
| | 09490 | hypothetical protein | 0 | 0 | No | 0 |
| | 09500 | hypothetical protein | 0 | 0 | No | 1 |
| II | 09510 | TaqI-like C-terminal specificity domain-containing protein | 0 | 0 | No | 2 |
| | 09520 | hypothetical protein | 0 | 0 | No | 0 |
| | 09530 | hypothetical protein | 0 | 0 | Yes | 1 |
| | 09540 | hypothetical protein | 0 | 1 | No | 1 |
| | 09550 | Protein of unknown function DUF4013 (IPR025098) | 0 | 6 | No | 1 |
| | 09560 | TetR regulator repressor; DNA binding; | 0 | 0 | No | 2 |
| | 09570 | hypothetical protein | 0 | 0 | No | 1 |
| | 09580 | SprT family zinc-dependent metalloprotease | 0 | 0 | No | 1 |
| | 09590 | hypothetical protein | 2 | 2 | No | 0 |
| | 13350 | ABC transporter, permease protein, BtuC-like (IPR000522) | 10 | 9 | Yes | 0 |
| | 13360 | iron ABC transporter permease (siderrophore like, Fe3+ transport) | 0 | 9 | No | 1 |
| | 13370 | ABC transporter substrate-binding protein B12 /periplasmic | 0 | 0 | Yes | 0 |
| | 13380 | ABC transporter substrate-binding protein | 0 | 0 | Yes | 0 |
| | 13390 | hypothetical protein (PKD, Ig, cohesin, CARB-binding domains) | 0 | 9 | Yes | 25 |
| | 13400 | hypothetical protein | 1 | 1 | Yes | 4 |
| | 13410 | hypothetical protein | 0 | 0 | Yes | 1 |
| | 13420 | hypothetical protein ( Ig, cohesin, CARDB/bt. cell adhesion, CARB-binding domains) | 0 | 0 | Yes | 26 |
| | 13430 | FCobN/magnesium chelatase | 0 | 0 | No | 6 |
| | 13440 | Magnesium-chelatase subunit ChlI-like (IPR045006) | 0 | 0 | No | 1 |
| | 13450 | hypothetical protein Mg-chelatase/VWA-domain | 0 | 0 | No | 1 |
| III | 13460 | nucleoside transporter/Fe-cotransport | 7 | 5 | No | 0 |
| | 13470 | ABC transporter ATP-binding protein/Fe-siderrophore | 0 | 0 | No | 3 |
| | 13480 | ABC transporter substrate-binding protein B12/periplasmic | 0 | 0 | Yes | 1 |
| | 13490 | Toxin-antitoxin system, RelE/ParE toxin family (proton donor motif) | 0 | 0 | No | 0 |
| | 13500 | hypothetical protein | 0 | 0 | No | 1 |
| | t00200 | tRNA-Ser | - | - | - | - |
| | 13510 | hypothetical protein | 0 | 0 | No | 1 |
| | 13520 | hypothetical protein (1TM, large cytoplasmic domain) | 1 | 1 | No | 0 |
| | 13530 | DEAD/DEAH box helicase (RNA helicase) | 0 | 0 | No | 1 |
| | 13540 | hypothetical protein (Tyr recombinase, CRE phage integrase, DNA break) | 0 | 0 | No | 2 |
| | 13550 | ABC transporter substrate-binding protein (MEMBRANE-BOUND LYTIC MUREIN TRANSGLYCOSYLASE F) | 0 | 0 | Yes | 0 |
| | 13560 | helix-turn-helix domain-containing protein | 0 | 0 | No | 1 |
| | 13570 | Uncharacterised conserved protein UCP006404, metalloprotease M50/CBS (IPR016483) | 0 | 9 | No | 1 |
| IV | 13800 | hypothetical protein (NosD, Pectin lyase fold/virulence) | 0 | 0 | Yes | 39 |
| | 14430 | hypothetical protein | 0 | 0 | No | 2 |
| | 14440 | glycosyltransferase family 4 protein | 0 | 0 | No | 2 |
| | 14450 | glycosyltransferase family 1 protein | 0 | 0 | No | 3 |
| | 14460 | glycosyltransferase family 1 protein | 0 | 0 | No | 0 |
| | 14470 | DUF2206 domain-containing protein | 0 | 17 | No | 6 |
| | 14480 | glycosyltransferase family 2 protein | 0 | 0 | No | 3 |
| | 14490 | flippase | 14 | 14 | No | 2 |
| | 14500 | dTDP-4-dehydrorhamnose reductase | 0 | 0 | No | 2 |
| | 14510 | dTDP-glucose 4,6-dehydratase | 0 | 0 | No | 1 |
| | 14520 | glucose-1-phosphate thymidylyltransferase RfbA | 0 | 0 | No | 3 |
| | 14530 | dTDP-4-dehydrorhamnose 3,5-epimerase | 0 | 0 | No | 1 |
| | 14540 | hypothetical protein (Ig-like fold) | 0 | 0 | Yes | 3 |
| | 14550 | hypothetical protein | 0 | 0 | No | 5 |
| | 14560 | hypothetical protein | 0 | 2 | No | 5 |
| | 14570 | hypothetical protein | 3 | 2 | No | 0 |
| V | 14580 | hypothetical protein | 0 | 0 | No | 3 |
| | 14590 | Ig-like domain repeat protein | 0 | 0 | No | 16 |
| | 14600 | DUF4350 domain-containing protein | 0 | 0 | No | 3 |
| | 14610 | MoxR family ATPase | 0 | 0 | No | 0 |
| | 14620 | DUF58 domain-containing protein | 0 | 0 | No | 3 |
| | 14630 | hypothetical protein | 0 | 0 | No | 1 |
| | 14640 | hypothetical protein (UDP-Glc/GDP-Man, NAD(P)-binding) | 0 | 0 | No | 2 |
| | 14650 | acyl-CoA dehydratase activase | 0 | 0 | No | 0 |
| | 14660 | acetate uptake transporter | 6 | 6 | No | 2 |
| | 14670 | hypothetical protein | 0 | 3 | No | 1 |
| | 14680 | double-cubane-cluster-containing anaerobic reductase | 0 | 0 | No | 0 |
| | 14690 | phosphate ABC transporter substrate-binding protein | 0 | 0 | Yes | 1 |
| | 14700 | hypothetical protein | 0 | 0 | Yes | 1 |
| | 14710 | TIGR00289 family protein | 0 | 0 | No | 0 |
| | 14720 | hypothetical protein | 1 | 1 | No | 3 |
| | t00230 | tRNA-Val | - | - | - | - |
| | 14730 | acetyl-CoA carboxylase biotin carboxylase subunit | 0 | 0 | No | 1 |
| | 16190 | hypothetical protein | 0 | 0 | No | 0 |
| | 16200 | hypothetical protein (DUF2202, Ferritin-like) | 0 | 1 | No | 2 |
| | 16210 | hypothetical protein | 0 | 0 | No | 0 |
| VI | 16220 | hypothetical protein (NosD domain, Pectin lyase fold/virulence domain) | 0 | 0 | No | 26 |
| | 16230 | hypothetical protein (NosD domain, Pectin lyase fold/virulence domain) | 0 | 0 | Yes | 7 |
| | 16240 | hypothetical protein | 0 | 1 | Yes | 4 |
| | 18180 | TIGR03576 family pyridoxal phosphate-dependent enzyme | 0 | 0 | No | 1 |
| | 18190 | DUF2098 domain-containing protein | 0 | 0 | No | 0 |
| | 18200 | carbonic anhydrase | 0 | 1 | Yes | 2 |
| | 18210 | twin-arginine translocase | 1 | 1 | Yes | 0 |
| | 18220 | twin-arginine translocase subunit TatC | 6 | 5 | No | 0 |
| | 18230 | hydrogenase maturation protease | 0 | 0 | No | 0 |
| | **18240** | **Ni/Fe hydrogenase large subunit** | 0 | 0 | No | 3 |
| VII | **18250** | **Ni/Fe hydrogenase small subunit (TAT signal, iron-sulfur cluster binding)** | 0 | 0 | No | 6 |
| | 18260 | hypothetical protein | 0 | 1 | Yes | 7 |
| | 18270 | hypothetical protein (FxLYD domain) | 1 | 1 | Yes | 2 |
| | 18280 | hypothetical protein (PIN-like domain) | 0 | 0 | No | 0 |
| | 18290 | hypothetical protein | 1 | 1 | No | 0 |
| | 18300 | endonuclease III | 0 | 0 | No | 1 |
| | 18310 | PD-(D/E)XK nuclease family protein | 0 | 0 | No | 0 |
| | 18320 | hypothetical protein (GTPase activity GTP binding) | 0 | 0 | No | 0 |

**Supplementary Table S2. List of genes located in the corrosive strain-specific gene islands.** Position numbers are the same as in Fig. 2c. InterPro and NetNGlyc 1.0 were used for transmembrane helix and signal peptide, and N-linked glycosylation site, respectively.

**Supplementary Table S3. Number of potential N-linked glycosylation sites in MIC hydrogenases and cell-surface related proteins in selected corrosive or non-corrosive *M. maripaludis* strains.** N-linked glycosylation sites were detected using NetNGlyc 1.0 (ref).

| Protein | Gene number in Mic1c10 | Number of NxS/T motif | | | | | | |
|---|---|---|---|---|---|---|---|---|
| | | Mic1c10* | OS7* | MIC098Bin5* | KA1* | JJ | S2 | C5 |
| AglB | 03010 | 3 | 3 | 3 | 3 | 3 | 3 | 4 |
| S-layer protein | 09120 | 2 | 2 | 2 | 2 | 3 | 1 | 0 |
| | 12420 | 1 | 1 | 1 | 1 | 0 | 0 | 0 |
| | 14270 | 3 | 0 | 3 | 3 | 2 | 0 | 0 |
| COG1361S-layer family protein | 07040 | 4 | 4 | 4 | 4 | 4 | 4 | 4 |
| | 11730 | 3 | 3 | 3 | 3 | 3 | 3 | 3 |
| | 12930 | 2 | 2 | 2 | 2 | 0 | 2 | - |
| cell wall-binding repeat-containing protein | 10740 | 2 | 3 | 3 | 3 | 3 | 3 | - |
| | 10750 | 16 | 15 | 16 | 16 | 17 | 16 | 14 |
| MIC island H2ase LSU [NiFe] | 18240 | 3 | 3 | 3 | 3 | - | - | - |
| MIC island H2ase SSU [Fe-S] | 18250 | 6 | 2 | 2 | 2 | - | - | - |

*Corrosive strains.

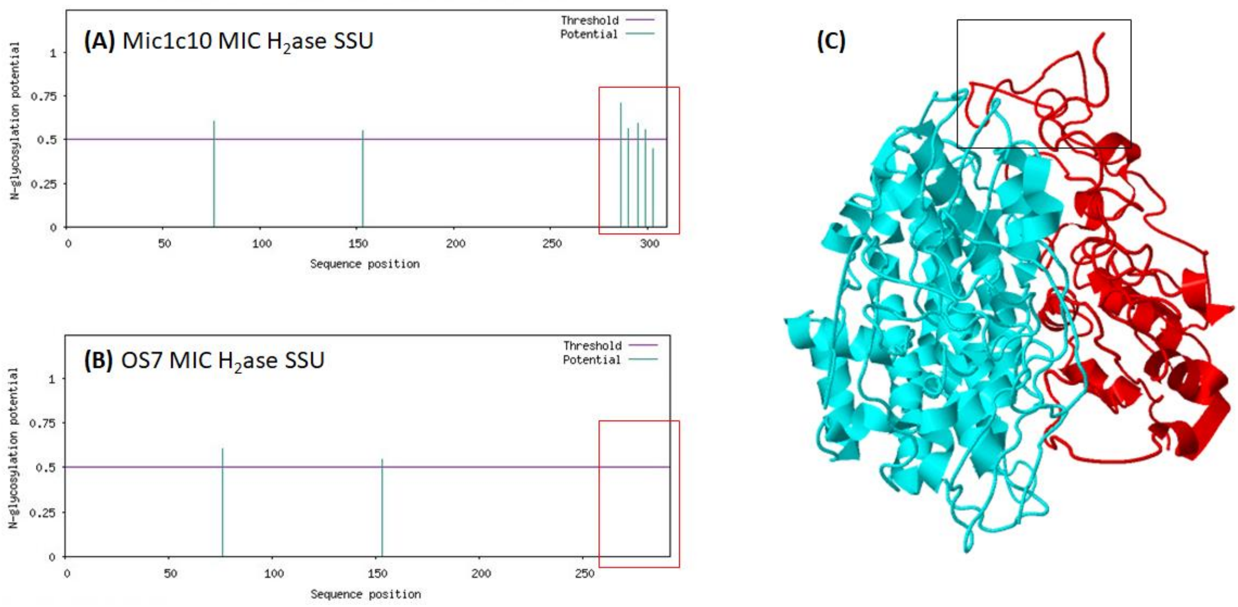


**Supplementary Figure 3. Potential N-linked glycosylation sites in the small subunit (SSU) of the MIC hydrogenases of strains Mic1c10 (A) and OS7 (B), and the predicted dimer structure of the MIC hydrogenase of strain Mic1c10.** The C-terminal end of the SSU, where four N-linked glycosylation sites unique to Mic1c10 are located, is highlighted. NetNGlyc 1.0 (ref) was used to detect the potential N-linked glycosylation sites, and PEPPI (ref) was used to predict the interaction between the two subunits of the MIC hydrogenase of Mic1c10.

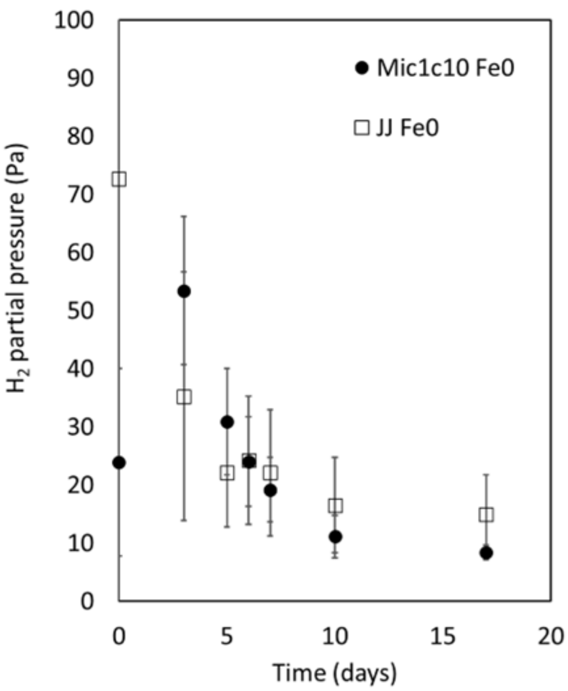


**Supplementary Figure 4. Change in the $H_2$ partial pressure (Pa) over time (days) for two different strains - corrosive Mic1c10 and non-corrosive JJ. The strains were grown on $Fe^0$, and the microbial cells were incubated as described in the text.**

Strain JJ (DSM 2067) was routinely cultivated in the modified NBRC medium 927 without $Fe^0$-granules but with $H_2/CO_2$ (80/20) as the electron and carbon sources. The experiment was initiated by inoculating 10% (v/v) culture of each strain to the modified NBRC medium 927 (5g $Fe^0$-granule in 50 mL medium). The pressure within each vial was monitored using a differential pressure manometer (RS-8890G, RS PRO), while the gas content was measured using the GC equipped with TCD. The incubation and measurements were extended until the vial got over-pressurized, indicating that the rate of $H_2$ evolution surpassed $H_2$ consumption. The experiment was conducted in triplicate, with the final data point for Mic1c10-$Fe^0$ being derived from a duplicate measurement.
